# Multisensory Continuous Psychophysics: Perceived Visual Object Location is Improved by Auditory Cues

**DOI:** 10.64898/2026.06.03.729954

**Authors:** Björn Jörges, Jong-Jin Kim, Laurence R. Harris

## Abstract

Continuous Psychophysics, which couples a continuous stimulus with a continuous response, is a promising tool to break out of the confines of traditional designs based on discrete trials. In this pre-registered study, we explore to what extent this paradigm is useful in the study of multisensory integration. We expand on Tonelli et al.’s (2025) seminal study by additionally examining the role of eye-movements, using a Kalman filter to estimate the sensory noise underlying behavioral tracking parameters and employing a virtual reality set-up. We immersed two cohorts of participants (n = 30 each) in a virtual meadow environment and asked them to continuously track a drone (Experiment 1) or a swarm of flies (Experiment 2) with a controller, while simultaneously recording their eye movements. We manipulated the reliability of visual cues using four levels of fog (from a completely clear view to impenetrable fog where no visual cues to the target’s position were available) as well as the presence of sound cues emitted from the object (sound present/absent). The maximum correlation between stimulus and response was higher when sound was present in some conditions, particularly when visual uncertainty was high, while the tracking delay remained unaffected across all fog levels. Using a Kalman filter to estimate the underlying sensory noise, we found strong evidence that sensory noise was lower when sound was present than when sound was absent both for manual and for ocular tracking, particularly for those conditions with higher visual uncertainty. In exploratory analyses, we further show strong correlations between manual and ocular tracking in all measures (maximum correlation, tracking delay, sensory precision). However, when isolating the multisensory advantage, these correlations all but disappeared for maximum correlation and tracking delay, while remaining substantial for sensory precision. Similarly, behavioral tracking correlated generally strongly with underlying sensory noise, but much less so when it came to the advantage conferred by added sound cues. Our results show that continuous psychophysics is well-suited for the study of multisensory integration, particularly when a Kalman filter analysis is used to estimate sensory uncertainty from behavioral data.

## Introduction

Most current models and conceptualizations of multisensory integration (Chandrasekaran, 2017; Drugowitsch et al., 2014; Ernst & Banks, 2002; Noppeney, 2021) are derived from trial-based experimental designs. Such models are restricted to instances of discrete decisions, such as locating the source of a single sound or estimating the direction in which an observer moves at some particular moment in time. However, real-life multisensory decision-making is often a continuous process of perception-and-action coupling, for example when we are tracking the siren of a moving ambulance over an extended period of time or following it with a car.

Many models of multisensory integration are based on causal inference (Noppeney, 2021): the organism first determines how likely two (or more) sensory stimuli are to originate from the same source. If a common source is unlikely, *segregation* occurs, and the two sensory stimuli are not integrated. Only when a common source is deemed likely are the sensory inputs integrated. Integration is commonly thought to be weighted as a function of the sensory uncertainty of each of the unimodal cues: maximum likelihood estimation (Ernst & Banks, 2002; Ernst & Bülthoff, 2004). As a result, the final multimodal percept is located between the unimodal cues and is represented with higher precision than any of the contributing sources.

In this project, we aim to confirm these principles using Continuous Psychophysics. Continuous Psychophysics (Bonnen et al., 2015; Décima et al., 2022; Falconbridge et al., 2023b, 2023a; Grillini et al., 2022; Jörges et al., 2024; Straub & Rothkopf, 2022) refers to a paradigm that couples a continuous stimulus with a continuous response, such as tracking a dot with a finger (Bonnen et al., 2015) or a cursor (Décima et al., 2022), nulling self-motion (Falconbridge et al., 2023b, 2023a), or pointing a joystick in the direction of self-motion (Jörges et al., 2024). While data are typically analyzed using a Kalman filter approach that allows an estimation of the sensory uncertainty of the transduced stimulus, other methods based on summary statistics of the responses have been proposed. Rather than assessing the underlying sensory uncertainty, these methods focus on behavioral tracking parameters like the tracking delay or the peak correlation between stimulus and response. These parameters typically correlate highly with the sensory uncertainty (e.g., Bonnen et al., 2015) and have therefore been proposed as easier-to-compute proxies for the precision of the sensory representation. However, this tight link between tracking delay, peak correlation, and sensory uncertainty is not a given *a priori* and might vary with stimulus or task characteristics. We address in an exploratory analysis whether this relationship holds for multisensory integration.

In a recent study, Tonelli et al. (2025) employed a continuous tracking design to assess audiovisual integration. In a vision-only condition, a visual stimulus was presented in 2D on a screen and moved horizontally in a random walk. In a sound-only condition, an audio target’s motion was simulated in the same way and presented using an array of loudspeakers. In a combined audio-visual condition, both the visual and the auditory components were presented simultaneously. The authors also included audiovisual conditions where the visual and the audio targets were shifted relative to each other in time such that either the visual target led the auditory target or the other way around. They did not find the expected increase in the correlation between stimulus and response in the bimodal condition, nor did they find improvements in the tracking delay over the unimodal conditions when both cues were present. Both these things would be expected in a causal inference framework, as multisensory integration should decrease the sensory noise. Decreases in sensory noise have, in turn, been shown to strongly correlate with improvements in these two tracking parameters in previous continuous psychophysics studies (e.g., [Bonnen et al., 2015; Jörges et al., 2024a]). One of the goals of the present study is, therefore, to estimate sensory uncertainty using a Kalman filter and to determine whether the absence of a multisensory advantage in tracking performance is related to the absence of multisensory advantages in sensory noise.

Kalman filters are a class of Bayesian models that update an estimate of the state of the world based on a representation of the variability in the world and the sensory uncertainty of incoming information. When applied to the study of perception, a) the estimate of the state of the world might correspond to an estimate of the position of a stimulus, b) the variability of the world might relate to a representation of how much its position changes over time, and c) the sensory uncertainty would correspond to noise introduced by the human sensory apparatus. If variability in the stimulus is low and the sensory uncertainty is high, then the updated estimate of the stimulus at a later time (i.e., at *t+1*) will be close to the previous estimate of the stimulus at time *t*. That is, the estimate will barely be updated. Conversely, if the stimulus is known to be highly variable and sensory uncertainty is low, the percept at t+1 will be very close to the position as reported by the sensory apparatus, i.e., the estimate of the stimulus will be updated by a large margin. The variability in the stimulus can be determined by the experimental design, such that the updating fraction is uniquely determined by the sensory uncertainty. This system can thus be used to estimate sensory uncertainty. A decrease in sensory uncertainty in a multimodal context could be used by the organism to improve tracking performance, e.g., by updating the estimate of the stimulus more completely (as captured by the Kalman filter), by tracking a target more closely (as captured by the maximum correlation between stimulus and response) or by responding faster to changes in the stimulus (as captured by the tracking delay).

It is a further open question in the application of Continuous Psychophysics as to how different response modalities might interact. While, for example, both manual (Bonnen et al., 2015; Décima et al., 2022) and ocular tracking (Grillini et al., 2022) have been used as experimental response modalities, to our knowledge no studies have recorded both at the same time. A secondary goal for this study is, therefore, to explore the interactions between manual and concurrent ocular tracking, both generally and as it pertains to multisensory integration.

In two sister experiments with slight variations in the motion parameters of the target, we varied the directional cues (visual and auditory) of a moving target (a drone in Experiment 1 and a swarm of flies in Experiment 2) which participants continuously tracked by pointing at it with a controller, while at the same time recording their eye movements.

This study addresses the following pre-registered goals:

- Replicate and expand on Tonelli et al.’s (2025) exploration of the use of Continuous Psychophysics for multisensory applications using the example of multisensory object tracking in a VR setting.
- Confirm Optimal Integration accounts of multisensory integration in a Continuous Psychophysics task by showing that sensory precision is higher when auditory and visual cues are present, using a Kalman filter approach.

In exploratory analyses, we also:

- Investigated the relationship between sensory precision (as estimated using a Kalman filter) and behavioral tracking parameters.
- Assessed the relationship between manual and concurrent ocular tracking in Continuous Psychophysics broadly, as well as specifically for multisensory integration.

## EXPERIMENT 1

### Methods

#### Participants

We collected data from 30 participants from the York University undergrad research participant pool. In a previous Continuous Psychophysics experiment (Jörges et al., 2024), we found that even 10 participants yielded precise effect size estimates. Participants received course credit for their participation. Informed consent was obtained in writing, and the experiments were conducted in line with the Declaration of Helsinki. Ethics approval was granted by the York University Office for Research Ethics under the protocol number e2025-303. Data collection for this experiment occurred between October 2025 and March 2026.

#### Apparatus

The stimulus was presented using an Alienware laptop (16 GB RAM, an Intel Core i7-9750H CPU (2.60 GHz), and an NVIDIA GeForce RTX 2060) with a VIVE Pro EYE head-mounted device (with a field of view of 110°, a resolution of 1440 × 1600 per eye, a 90 Hz refresh rate and a built-in Tobii Eye Tracker). The stimulus was programmed in Unity 2021.3.13f1 and both the Unity project and the executable are available for download on Open Science Foundation (https://osf.io/6q7x3/). Data was analyzed in R (v.4.3.2). Manual responses were collected using a VIVE controller whose position and rotation was recorded at 90 Hz through the Unity program. Eye movements were recorded using the built-in eye tracker.

#### Stimulus

##### Visual stimulation

We presented participants with a drone flying left and right on an arc that ranged between -25 and +25° around the straight ahead of the observer at a distance of 12.5 m and at 2 m above the simulated ground. See Figure 1.

**Figure 1:**
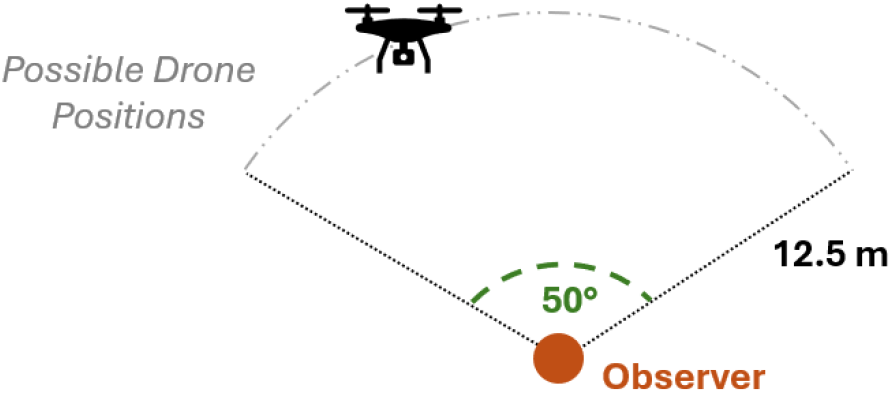
Diagram of the setup seen from a above.

The participant’s first-person viewpoint was simulated at an average adult sitting eye-height of 1.30 m above the ground. The drone’s position was determined by a random walk where each new angle of the drone relative to the observer was calculated by adding an angle drawn from a normal distribution with a mean of 0 and a standard deviation of 20°. The drone then spent the next 200 ms travelling from the position relative to the observer at t to the position at t+1 at a constant speed. The drone moved on an arc 12.5 m in front of the observer within +/-25° of the observer’s straight-ahead.

See Figure 1. We pre-generated nine trajectories and Figure 2 shows the simulated angle relative to the observer over the course of one of the 150 s runs. We simulated densities of fog (i.e., fog levels; NO FOG, MEDIUM FOG, HEAVY FOG, IMPENETRABLE FOG) to manipulate visibility and, with it, reliability of the visual cues, see Figure 3. In the IMPENETRABLE FOG condition, the drone was not visible to the observer. This fog level was only presented when sound cues were present.

**Figure 2.**
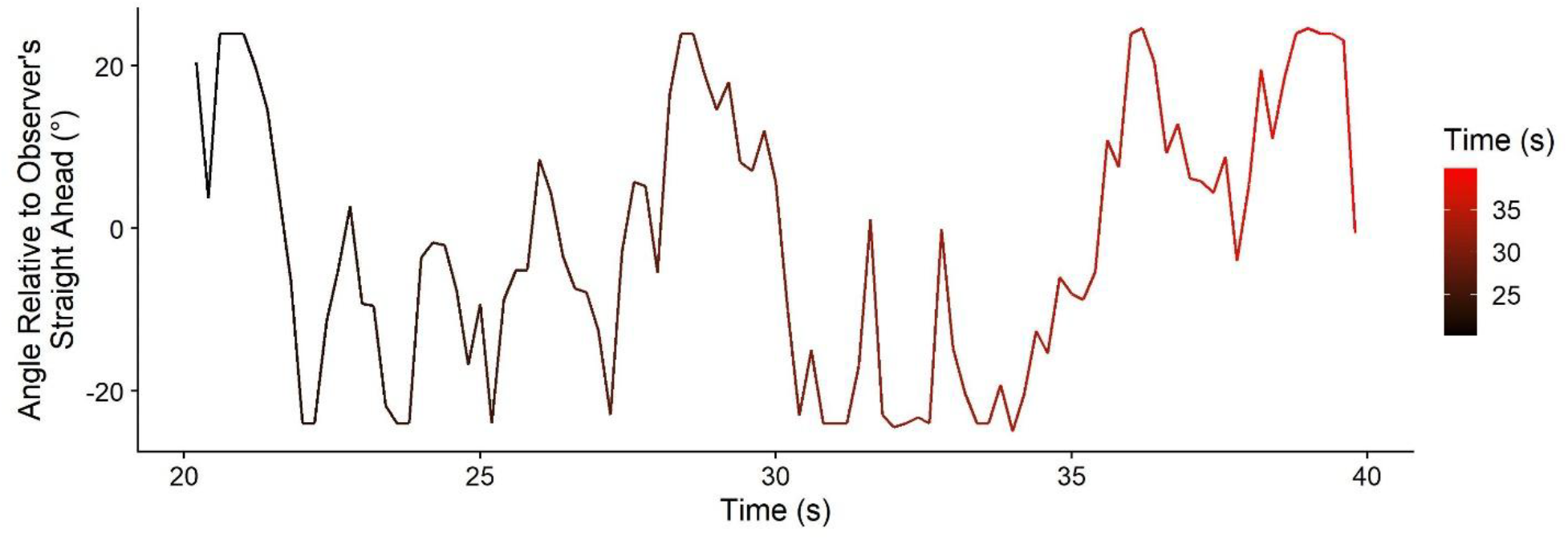
Angle relative to the observer (y axis) over the course of 20 s from one run (x axis). Note that, as part of the experimental design, we limited drone motion to +-25° relative to the observer’s straight-ahead.

**Figure 3.**
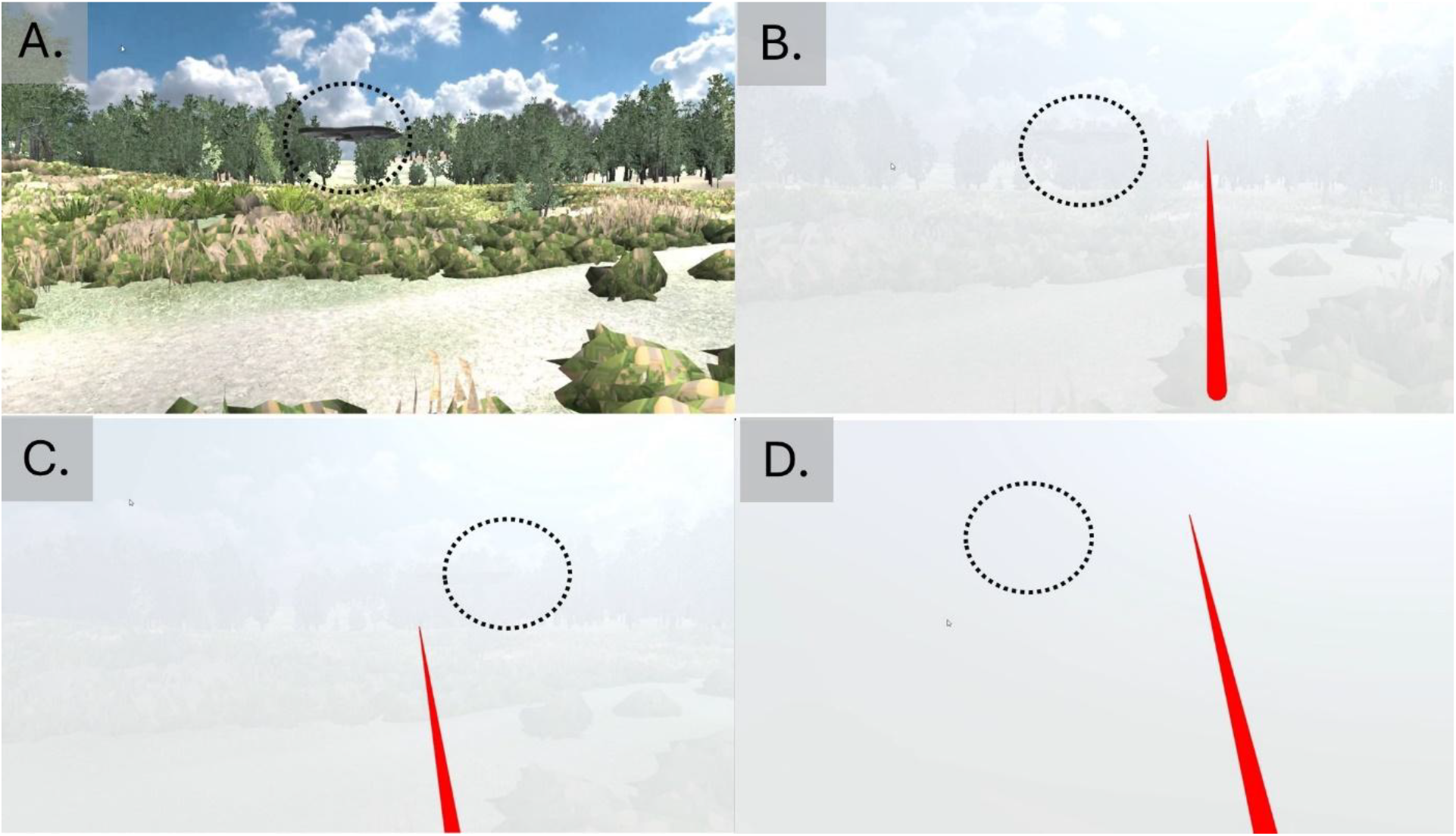
Screenshots from the experiment. A. NO FOG condition. B. MEDIUM FOG condition. C. HEAVY FOG condition. D. IMPENETRABLE FOG condition. The dotted circle was not present in the experiment but added in this figure to indicate the location of the drone. Videos of all conditions (including with and without sound) can be found in the Open Science Foundation repository (https://osf.io/6q7x3/).

##### Auditory stimulation

For auditory stimulation, we simulated a humming sound that was emitted from the drone. We used the built-in head transfer functions from Unity to spatialize the sound. Depending on the condition, sound was either present or absent, and it always emanated from the drone.

#### Procedure

Participants were first fitted with the headset and given the controller. The headset’s interpupillary distance was adjusted until participants confirmed a sharp and comfortable viewing experience. They were then given general instructions on the task: to continuously track the drones with a controller while also keeping their eyes on it. The controller was emitting a laser-like ray such that participants had feedback available on where they were pointing the controller. Then, we immersed them in a training run without fog until they confirmed that they were comfortable with the task. They then completed the seven test sessions (NO FOG + NO SOUND, MEDIUM FOG + NO SOUND, HEAVY FOG + NO SOUND, NO FOG + SOUND, MEDIUM FOG + SOUND, HEAVY FOG + SOUND, IMPENETRABLE FOG + SOUND) in randomized order. Each test session was 150s, and generally, it took less than 30 minutes to complete all sessions (including set up, instructions, training and main experimental conditions) and no breaks were needed.

#### Data Analysis

The pilot data as well as all code used for analysis can be downloaded from Open Science Foundation (https://osf.io/6q7x3/).

##### Pre-Processing

We first computed the position at which the laser emitted by the participant’s controller and/or their gaze, respectively, intersected the rounded plane on which the drone moved in space. Since the drone moved around the participant in a semicircle, we neglected variation in y direction and computed only the angle of the intersection between the laser and the rounded plane relative to the observer.

##### Cross-correlogram analysis

We then employed a cross-correlogram analysis to obtain estimates of how fast participants reacted to changes in the stimulus as well as how precisely they were able to do so, separately for the manual tracking data and the gaze tracking. Figure 4A shows the presented stimulus angle and the participant’s manual response angle from the first few seconds of a pilot run. We largely followed a procedure we have used before in a visual study (Jörges et al., 2024), which was adapted from the seminal paper by Bonnen et al. (2015). Cross-correlogram analyses compare how strong correlations between changes in the stimulus and changes in the participant responses are for different lags between stimulus and response, as shown in Figure 4B. The Maximum Correlation indicates precision in responses while the time lag at which this maximum correlation occurs (subsequently referred to as “Tracking Delay”) indicates how fast participants react to changes in the stimulus, i.e. the delay in tracking.

**Figure 4.**
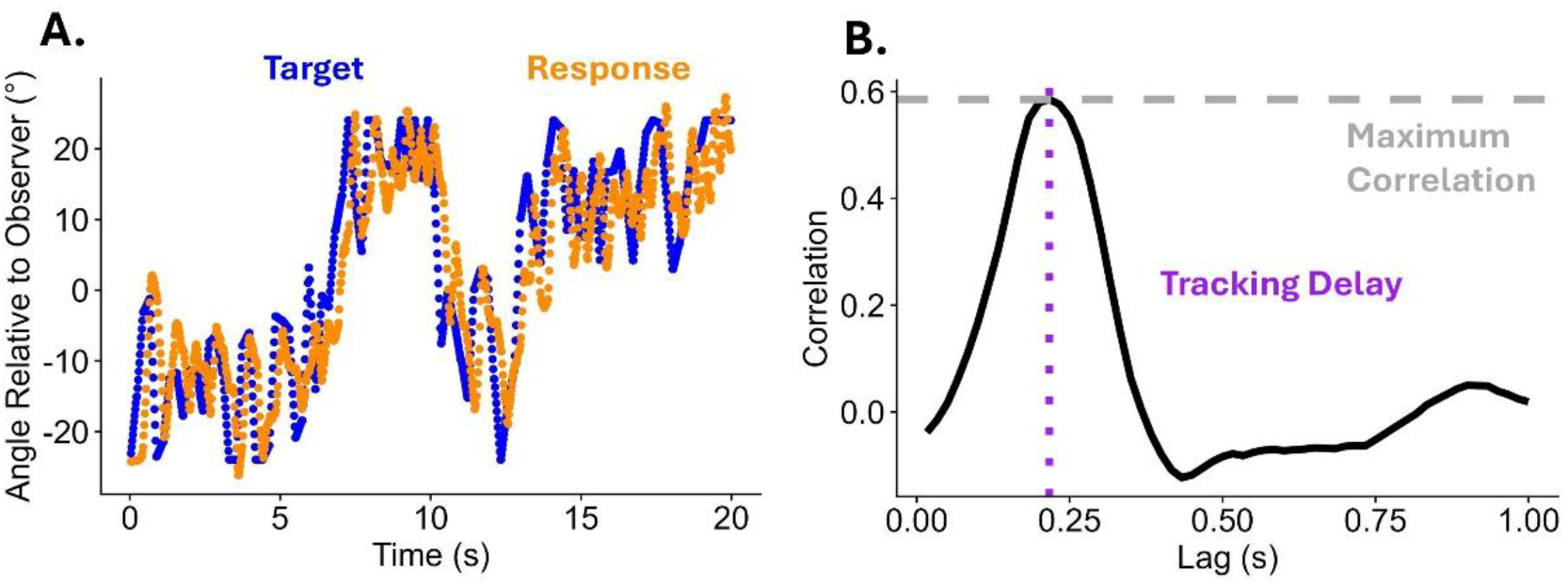
A. Target angle (blue) and manual response angle (orange) relative to the observer over the course of a partial run (x axis). Note that, as part of the experimental design, we limited drone motion (blue) to +-25° relative to the observer’s straight-ahead. B. Correlation between target angle and response angle (y axis) for different lags (x axis). The horizontal dashed line indicates the maximum correlation, and the purple dotted line indicates the tracking delay.

##### Outlier analysis

We excluded one run from one participant because no manual tracking data was recorded for parts of the trajectory due to equipment failure. After this exclusion, all runs from all participants passed the outlier criterion outlined in the pre-registration, i.e., no tracking delay at the bounds of the time lag interval ([0;1s]) was found for any of the runs.

##### Bootstrap to determine statistical significance

We then used the same bootstrap approach as in our previous study (Jörges et al., 2024) to determine statistically significant differences in maximum correlation and tracking delay between the test conditions. In this method, we create 1000 sub-samples from the whole dataset that each include 50% of the data points from 50% of the participants. For each of these sub-samples, we then fitted linear mixed models, separately for the dependent variables Manual Maximum Correlation, Gaze Maximum Correlation, and Time Lag of Manual Maximum Correlation, and Time Lag of Gaze Maximum Correlation using the R (R Core Team, 2017) package lme4 (Bates et al., 2015), with Condition (No Fog – No Sound, Medium Fog – No Sound, Heavy Fog – No Sound, No Fog – Sound, Medium Fog – Sound, Heavy Fog – Sound) as fixed effect and random intercepts per participant. In Wilkinson & Rogers (Wilkinson & Rogers, 1973) notation, this reads as:

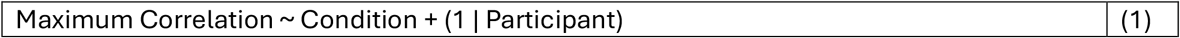

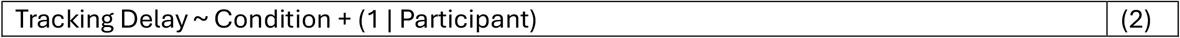

We then recorded the regression coefficients pertaining to the difference contrasts between sound present and absent within a certain fog level, i.e., No Fog – No Sound versus No Fog – Sound, Medium Fog – No Sound versus Medium Fog – Sound, Heavy Fog – No Sound versus Heavy Fog – Sound, for each of these sub-samples. We then took the 2.5% and 97.5% percentiles as lower and upper bound of the 95% confidence interval on these difference contrasts.

##### Predictions: Cross-correlations

Our prediction was that the maximum correlation (as an indicator of precision) would be higher in the multisensory conditions than in the corresponding unimodal (visual-only) conditions. For the tracking delay, we expected a decrease for the multisensory conditions as compared to the unimodal condition, indicating that participants adjusted their responses faster when sound cues were added.

##### Kalman filter

We further fitted a Kalman filter, a common method of analysis used in Continuous Psychophysics (Bonnen et al., 2015; Jörges et al., 2024), to obtain estimates of the sensory uncertainty associated with each experimental condition. Kalman filters are a subclass of Bayesian models that posit that observers update their responses to a larger or lesser degree (captured by the Kalman gain parameter *K*) depending on the sensory uncertainty and the variability in the stimulus as a prior. Given that we know how much participants update their responses and how large the variability in the stimulus is, we can then fit a parameter that captures the sensory uncertainty for each condition.

Specifically, the posterior variance (i.e., the variance in the participant responses: P) depends on the sensory uncertainty R and variance of the prior Q in the following way:

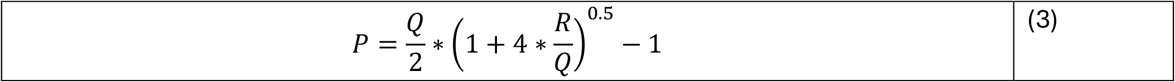

The Kalman gain K is then computed as:

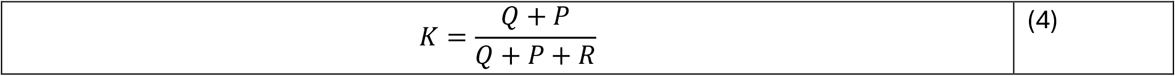

Having obtained the Kalman gain K, the participant response at t +1 () is estimated as participant response at t updated by a fraction (the Kalman gain) of the stimulus at t + 1. In line with pilot results that showed a fairly fast participant response to changes in the stimulus, we used a lag of 0.25s between participant response and stimulus.

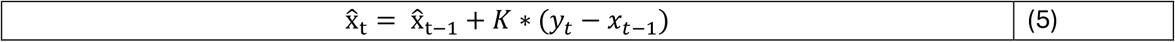

The residuals between predicted and observed performance is expected to follow a normal distribution with a mean of 0 and a standard deviation of K^2/R (Bonnen et al., 2015). Thus, we minimized the log likelihood for the residuals to follow a normal distribution with these parameters to fit the sensory uncertainty parameter R.

We then performed a statistical analysis over the fitted R parameters using a linear mixed model. We used R as dependent variable, the condition as fixed effect and random intercepts per participant as random effects.

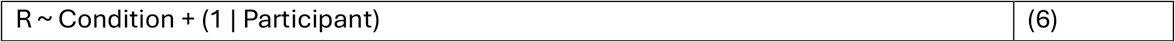

We then used the confint() function from base R to compute 95% confidence intervals on the difference contrasts of interest, i.e., No Fog – No Sound versus No Fog – Sound, Medium Fog – No Sound versus Medium Fog – Sound, Heavy Fog – No Sound versus Heavy Fog – Sound.

##### Predictions: Kalman filter

We expected sensory uncertainty to be lower for the multisensory conditions than the unimodal conditions.

## Results Experiment 1

The full results can be found in Table 1 (for the maximum correlation), Table 2 (for the tracking delay) and Table 3 (for the sensory noise parameters), and all data for this experiment (maximum correlation, tracking delay, sensory noise) are visualized in Figure 5.

**Table 1.** Experiment 1: 95% Confidence intervals for the difference contrasts for the Maximum Correlation, capturing performance for vision + sound in comparison to vision.

| **MAXIMUM CORRELATION** | Difference Contrast | 95% Confidence Interval (Lower Bound) | 95% Confidence Interval (Upper Bound) | Significance |
| --- | --- | --- | --- | --- |
| **Manual Tracking** |  |  |  |  |
| No Fog | -0.003 | -0.023 | 0.018 | n.s. |
| Medium Fog | -0.008 | -0.039 | 0.021 | n.s. |
| Heavy Fog | 0.019 | -0.017 | 0.051 | n.s. |
| **Ocular Tracking** |  |  |  |  |
| No Fog | 0.013 | -0.006 | 0.031 | n.s. |
| Medium Fog | -0.004 | -0.024 | 0.018 | n.s. |
| **Heavy Fog** | **0.023** | **0.004** | **0.039** | **\*** |

**Table 2.** Experiment 1: 95% Confidence intervals for the difference contrasts for the tracking delay, capturing performance for vision + sound in comparison to vision.

| **TRACKING DELAY** | Difference Contrast | 95% Confidence Interval (Lower Bound) | 95% Confidence Interval (Upper Bound) | Significance |
| --- | --- | --- | --- | --- |
| **Manual Tracking** |  |  |  |  |
| No Fog | -0.001s | -0.012s | 0.006s | n.s. |
| Medium Fog | 0.001s | -0.007s | 0.01s | n.s. |
| Heavy Fog | 0.001s | -0.013s | 0.013s | n.s. |
| **Ocular Tracking** |  |  |  |  |
| No Fog | -0.002s | -0.018s | 0.014s | n.s. |
| Medium Fog | 0.004s | -0.012s | 0.021s | n.s. |
| Heavy Fog | -0.001s | -0.028s | 0.021s | n.s. |

**Table 3.**
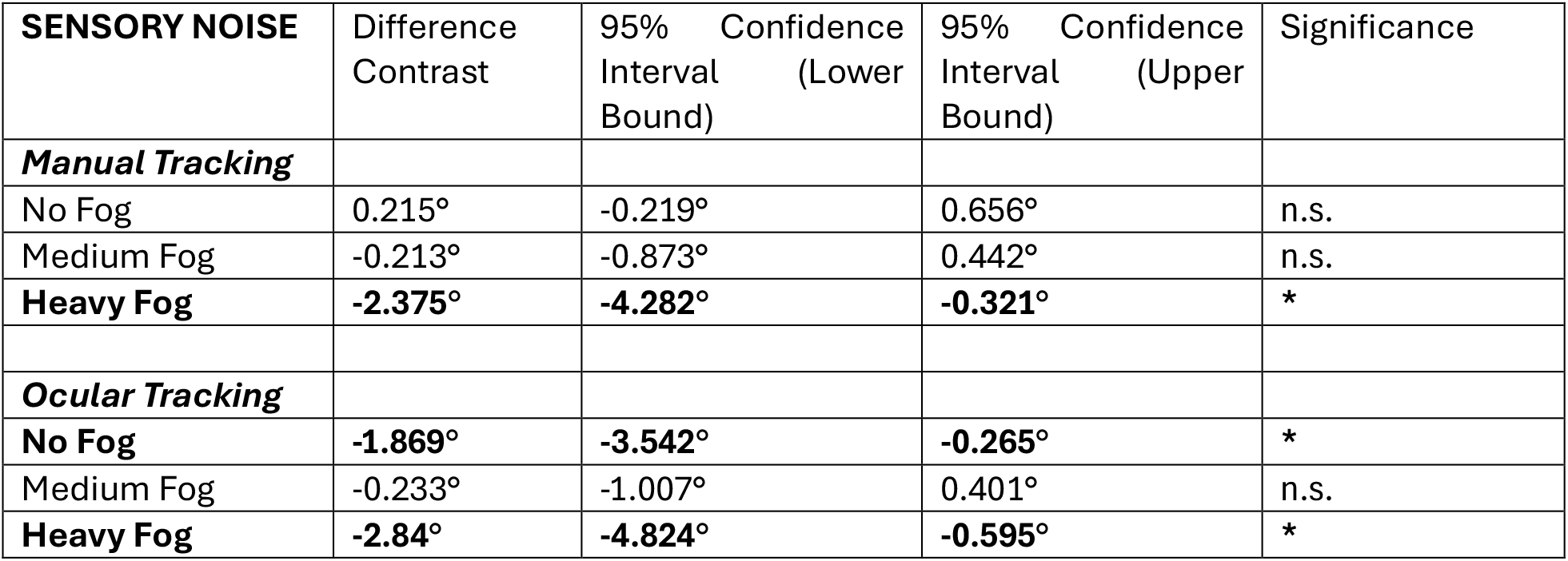
Experiment 1: 95% Confidence intervals for the difference contrasts for the Kalman filter gain, capturing performance for vision + sound in comparison to vision.

| SENSORY NOISE | Difference Contrast | 95% Confidence Interval (Lower Bound) | 95% Confidence Interval (Upper Bound) | Significance |
| --- | --- | --- | --- | --- |
| **Manual Tracking** |  |  |  |  |
| No Fog | 0.215° | -0.219° | 0.656° | n.s. |
| Medium Fog | -0.213° | -0.873° | 0.442° | n.s. |
| **Heavy Fog** | **-2.375°** | **-4.282°** | **-0.321°** | **\*** |
| **Ocular Tracking** |  |  |  |  |
| No Fog | **-1.869°** | **-3.542°** | **-0.265°** | **\*** |
| Medium Fog | -0.233° | -1.007° | 0.401° | n.s. |
| **Heavy Fog** | **-2.84°** | **-4.824°** | **-0.595°** | **\*** |

**Figure 5.**
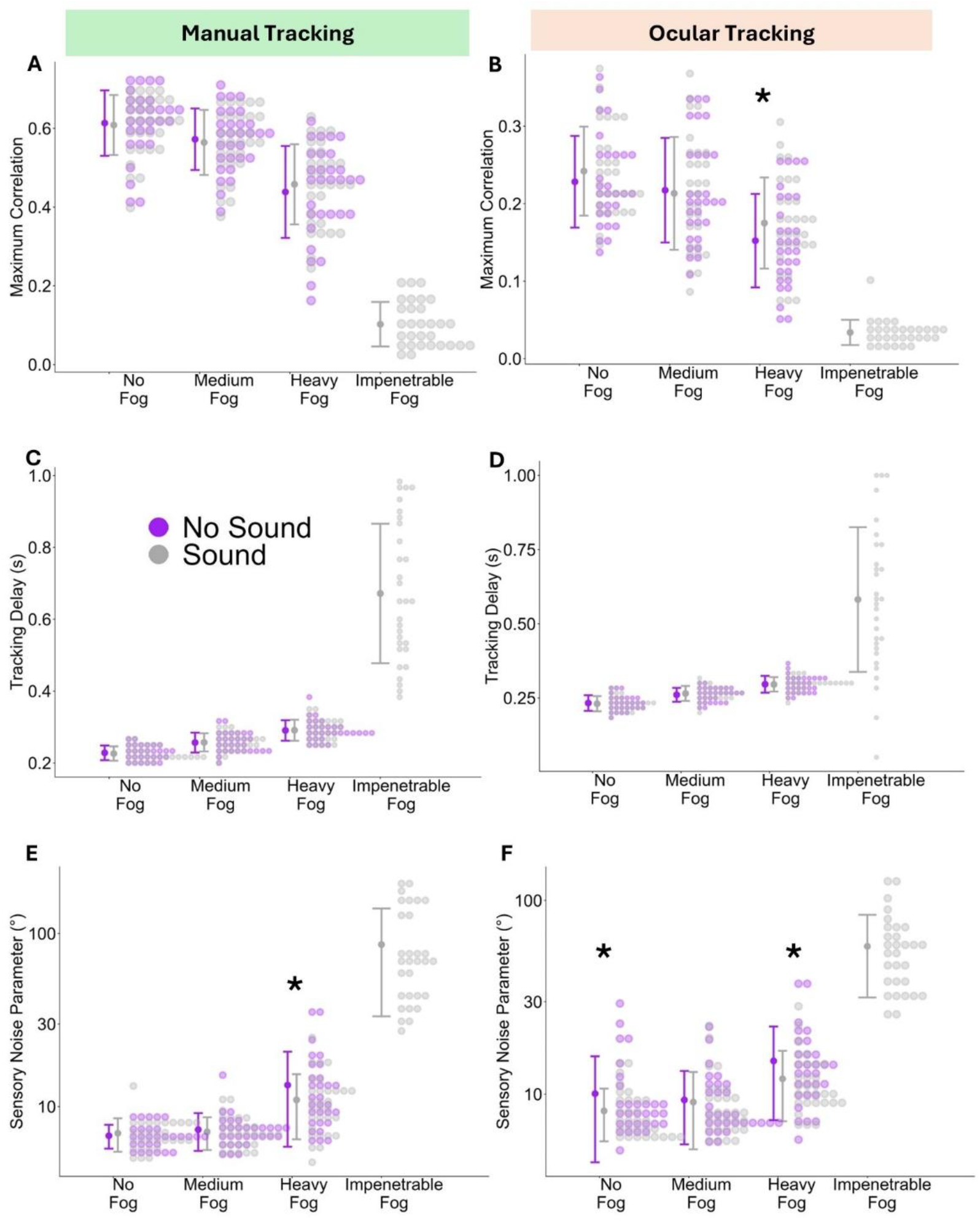
Experiment 1 – Maximum correlation (A, B), tracking delay (C, D) and sensory noise (E, F) for each fog level (x axis), separated by whether sound cues were present or absent (color-coded). The bold dots correspond to the mean for each condition and the antennae represent the standard deviation. The individual smaller dots represent the full distributions of data points. The left column (A, C, E) shows values for manual tracking and the right column (B, D, E) shows values for ocular tracking. The stars indicate where VISION + SOUND was significantly different from VISION ONLY.

Overall, the only dependent variable that showed a multisensory improvement for manual tracking was the Kalman filter sensory noise parameter in the HEAVY FOG condition (see Table 1 and Figure 5E). For ocular tracking, we found a significant increase in the maximum correlation for the HEAVY FOG condition (Table 1 and Figure 5B) and decreased sensory noise in the NO FOG and HEAVY FOG conditions (Table 3 and Figure 5F). Note that, while some trends in the expected direction are apparent in maximum correlations and sensory noise parameters even in the non-significant conditions, the 95% confidence intervals for the tracking delay are closely centered around 0 for all fog conditions and both manual and ocular tracking (Table 2 and Figure 5C and D).

## Discussion Experiment 1

The first experiment shows promising results for the use of Continuous Psychophysics in multisensory integration: while there were mixed results for behavioral tracking parameters (maximum correlation and tracking delay), multisensory cues increased sensory precision reliably in those conditions where visual information was least reliable. Based on these results, we devised a second experiment for three reasons: 1) we wanted to confirm the results regarding sensory precision, 2) we wanted to elucidate whether the mixed findings in the behavioral tracking parameters (maximum correlation and tracking delay) were due to lacking statistical power or the absence of a true effect, and 3) we wanted to address a small violation of the assumptions of the Kalman filter that we had committed in the first experiment with the intention of increasing ecological validity: the Kalman filter assumes that the position of the stimulus changes according to a random walk on each frame. In the first experiment, a new position for the drone was chosen every 200 ms, and the drone moved from its old position to the new position with a constant speed over this period. This made for a smoother, more natural motion, but it may have allowed participants to not only use positional information, but also speed information, which is neglected by the Kalman filter. We therefore designed a variation in which participants were asked to track a swarm of flies instead of a drone. The fuzzy nature of this tracking target allowed us to change its position frame-by-frame without breaking participants’ immersion while preventing them from using speed information.

## EXPERIMENT 2

### Participants

30 new participants completed this experiment between November 2025 and March 2026. They matched the participants from Experiment 1 to the extent described in the manuscript.

### Stimulus and Procedure

We used a swarm of flies as the tracking target instead of the drone (see Figure 6). It was generated using the Unity particle system such that the density of fly particles was highest in the center and tapered off towards the edges in a roughly Gaussian fashion. This mirrors the Gaussian blobs used in 2D Continuous Psychophysics experiments (Bonnen et al., 2015; Décima et al., 2022; Tonelli et al., 2025). Its motion was a true random walk where a new position was chosen for the swarm of flies each frame by choosing a value from a normal distribution with a mean of 0° and a standard deviation of 2° and adding it to its current position. See Figure 7 for the angles at which the swarm was represented relative to the observer over the course of one run. We also changed the drone buzzing emanating from the drone in Experiment 1 to a more appropriate insect-like buzzing. We further adjusted the density of the fog to qualitatively match the task difficulty from Experiment 2 for the Medium Fog and Heavy Fog conditions.

**Figure 6:**
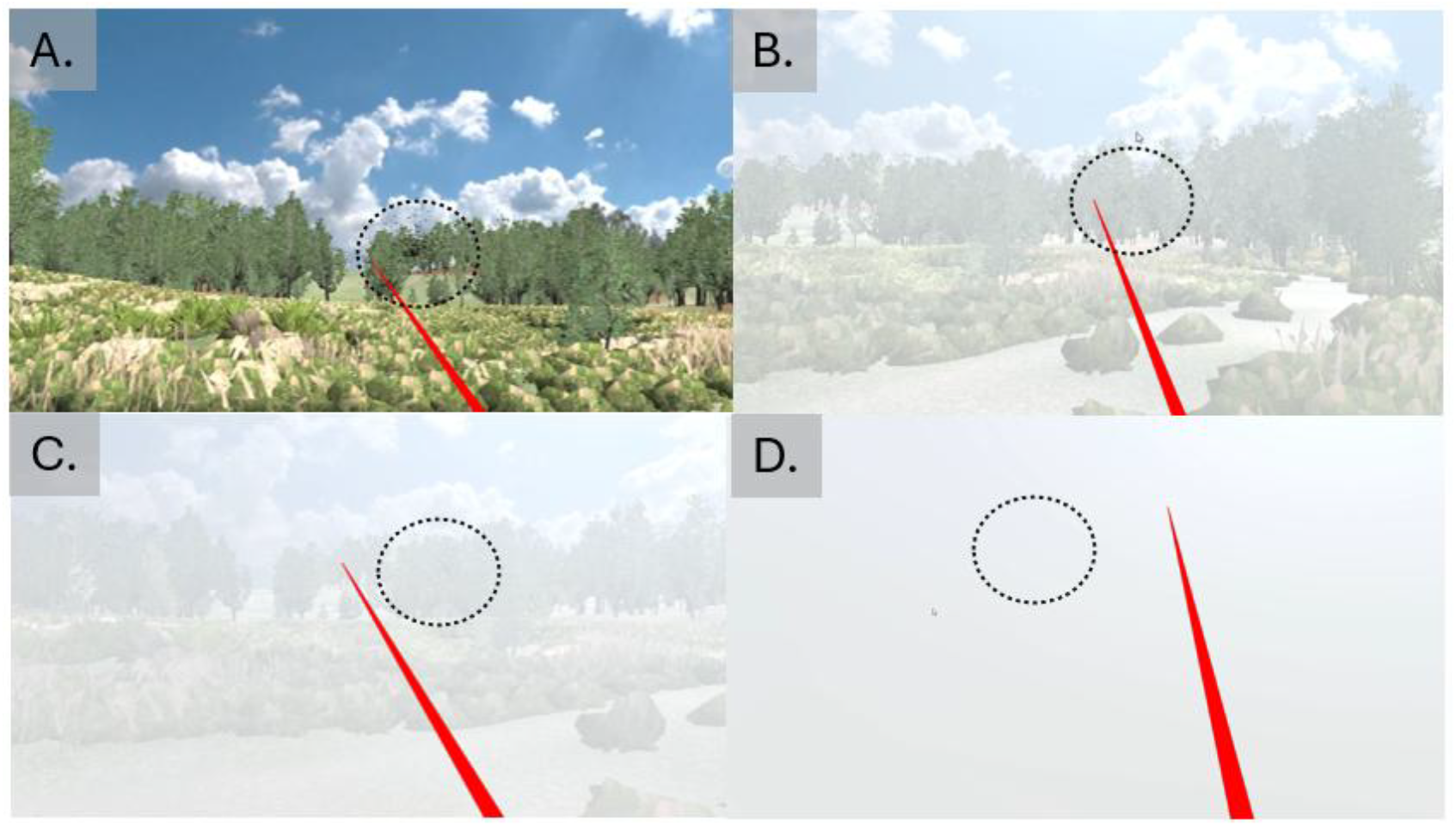
Screenshots from Experiments 2. A. No Fog condition. B. Medium Fog condition. C. Heavy Fog condition. D. Impenetrable Fog condition. The dotted circle was not present in the experiment, but added in this figure to indicate the location of the insect swarm. Videos from all conditions can be found in the Open Science Foundation repository (https://osf.io/6q7x3/).).

**Figure 7.**
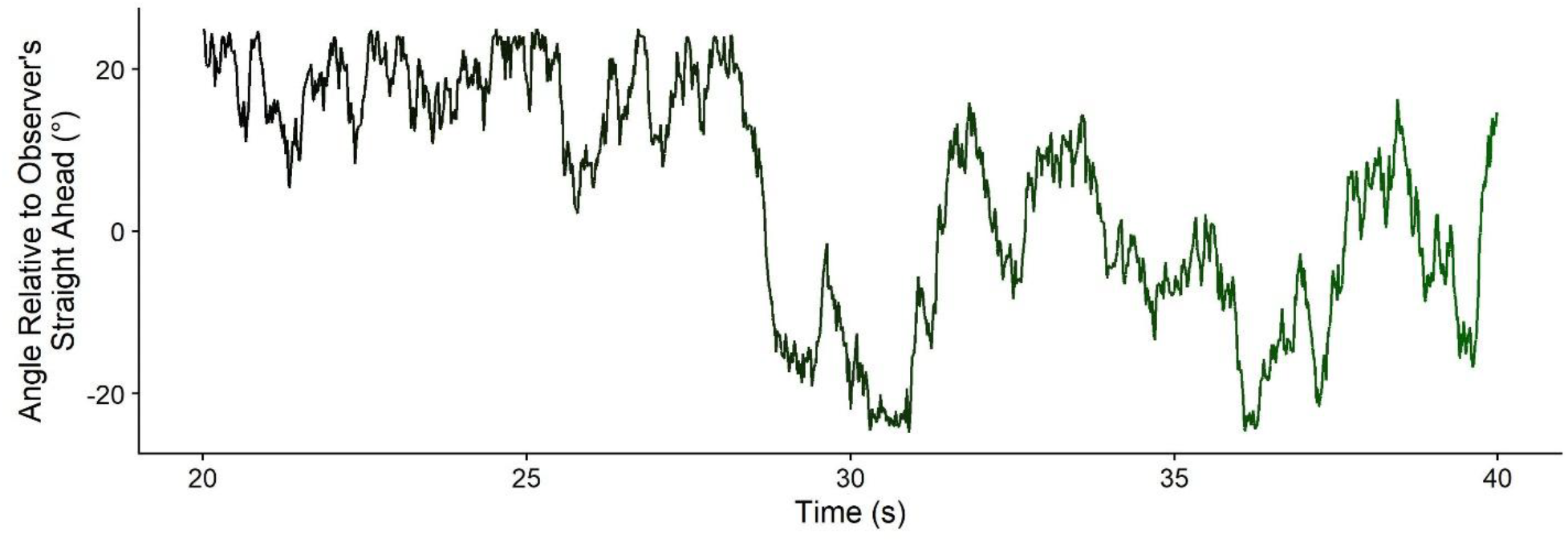
Angle of the swarm of flies relative to the observer’s straight ahead over 20s from one run. Note that, as part of the experimental design, we limited the swarm motion to +-25° relative to the observer’s straight-ahead.

### Data Analysis

We used the same Data Analysis procedure as for Experiment 1. In this experiment, we excluded three runs from one participant because the equipment failed to record manual tracking data, and one run each from two participants because no eye-tracking data was recorded for parts of the run. All other runs from all participants passed the outlier criterion and no further exclusions were necessary.

## Results Experiment 2

For the swarm experiment (i.e., Experiment 2), the full difference contrasts and 95% confidence intervals can be found in Table 4 (maximum correlation), Table 5 (tracking delay), and Table 6 (sensory noise), and Figure 8 visualizes the data.

**Table 4.** Experiment 2: 95% Confidence intervals for the difference contrasts for the Maximum Correlation, capturing performance for vision + sound in comparison to vision.

| **MAXIMUM CORRELATION** | Difference Contrast | 95% Confidence Interval (Lower Bound) | 95% Confidence Interval (Upper Bound) | Significance |
| --- | --- | --- | --- | --- |
| **Manual Tracking** |  |  |  |  |
| No Fog | 0.006 | -0.007 | 0.019 | n.s. |
| Medium Fog | 0.009 | -0.005 | 0.022 | n.s. |
| **Heavy Fog** | **0.01** | **0.002** | **0.017** | **\*** |
| **Ocular Tracking** |  |  |  |  |
| No Fog | 0.009 | -0.001 | 0.015 | n.s. |
| Medium Fog | -0.003 | -0.008 | 0.005 | n.s. |
| Heavy Fog | 0.003 | -0.002 | 0.009 | n.s. |

**Table 5.** Experiment 2: 95% Confidence intervals for the relevant difference contrasts for the tracking delay, capturing performance for vision + sound in comparison to vision.

| **TRACKING DELAY** | Difference Contrast | 95% Confidence Interval (Lower Bound) | 95% Confidence Interval (Upper Bound) | Significance |
| --- | --- | --- | --- | --- |
| **Manual Tracking** |  |  |  |  |
| No Fog | -0.006s | -0.021s | 0.001s | n.s. |
| Medium Fog | 0.003s | -0.052s | 0.041s | n.s. |
| Heavy Fog | -0.041s | -0.123s | 0.092s | n.s. |
| **Ocular Tracking** |  |  |  |  |
| No Fog | -0.02s | -0.101s | 0.044s | n.s. |
| Medium Fog | -0.004s | -0.088s | 0.081s | n.s. |
| Heavy Fog | 0.027s | -0.118s | 0.179s | n.s. |

**Table 6.** Experiment 2: 95% Confidence intervals for the relevant difference contrasts for the Kalman filter gain, capturing performance for vision + sound in comparison to vision.

| **SENSORY NOISE** | Difference Contrast | 95% Confidence Interval (Lower Bound) | 95% Confidence Interval (Upper Bound) | Significance |
| --- | --- | --- | --- | --- |
| **Manual Tracking** |  |  |  |  |
| No Fog | -0.27° | -1.113° | 0.56° | n.s. |
| **Medium Fog** | **-4.21°** | **-6.728°** | **-1.964°** | **\*** |
| **Heavy Fog** | **-8.502°** | **-12.833°** | **-4.054°** | **\*** |
| **Ocular Tracking** |  |  |  |  |
| No Fog | -2.438° | -4.161° | -0.619° | * |
| Medium Fog | -3.017° | -4.844° | -0.949° | * |
| Heavy Fog | -5.393° | -9.245° | -1.091° | * |

**Figure 8.**
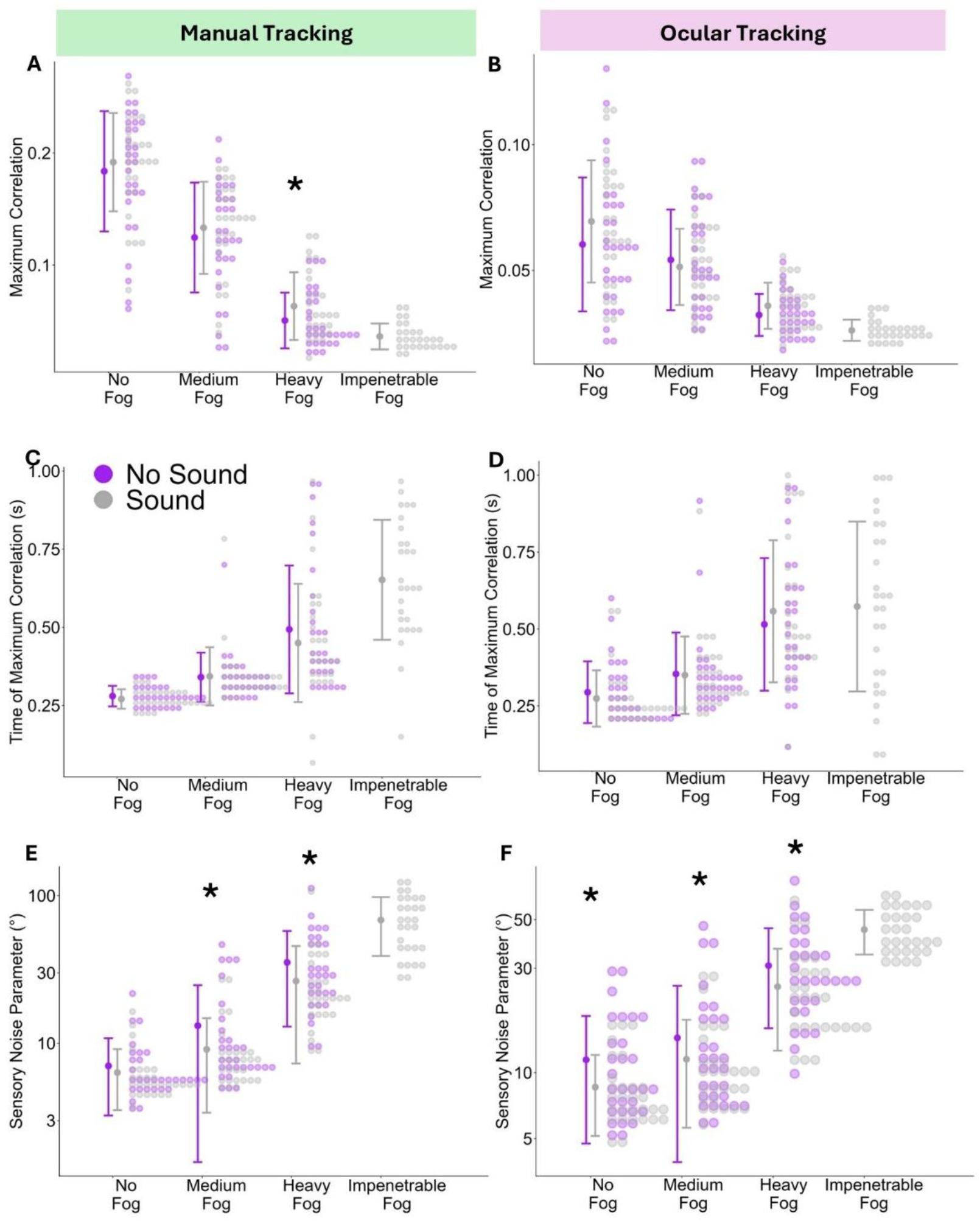
Experiment 2 – maximum correlation (A, B), tracking delay (C, D) and sensory noise (E, F) for each fog level (x axis), separated by whether sound cues were present or not (color-coded). The bolt dots correspond to the mean for each condition and the antennae represent the standard deviation. The individual smaller dots represent the full distributions of data points. The left column (A, C, E) shows values for manual tracking and the right column (B, D, E) shows values for ocular tracking. The stars indicate where VISION + SOUND was significantly different from VISION ONLY.

In terms of the maximum correlation, we only found a significant multisensory advantage for manual tracking in the HEAVY FOG condition (Table 4 and Figure 8A). As for Experiment 1, the tracking delay was not affected significantly by adding sound cues to the visual stimulus (Table 5 and Figure 8C and D).

For the sensory noise, we found multisensory improvements for manual tracking in the MEDIUM FOG and HEAVY FOG conditions (Table 6 and Figure 8E), and sensory noise derived from ocular tracking was decreased significantly in all three fog conditions (Table 6 and Figure 8F).

## Interim Discussion

The pattern in results for Experiment 2 was similar to those of Experiment 1: we found mixed results for a multisensory advantage in the maximum correlation between stimulus and response, and convincing evidence for an increase in sensory precision via the Kalman filter gains. As for Experiment 1, the tracking delay was unaffected by the addition of sound cues.

While this provides a clearer picture regarding the main hypothesis of this project (as in the pre-registered version), this rich data set provides an opportunity to shine light on two other questions: first, as outlined in the introduction, sensory precision tends to correlate highly with tracking delay and the maximum correlation. Given the discrepancy in results between behavioral tracking parameters (tracking delay and maximum correlation) on one hand and sensory precision on the other, we wanted to explore whether multisensory advantages in the latter were predictive of the former. Further, while manual and ocular tracking have been used separately in Continuous Psychophysics, we are, to our knowledge, the first to collect manual and ocular tracking data simultaneously. In this exploratory analysis, we will therefore study the link between manual and ocular tracking in general, as well as their relationship with multisensory integration.

## Exploratory Analysis: Sensory Noise, Tracking Delay and Maximum Correlation

In the seminal study on Continuous Psychophysics (Bonnen et al., 2015), the authors found strong correlations between behavioral tracking parameters (maximum correlation and tracking delay) and the sensory noise parameters estimated using a Kalman filter. In our exploratory analysis, we examine whether this relationship holds true in the present dataset, and how any such correlation relates specifically to performance advantages that might be attributable to multisensory integration.

### How well does sensory precision correlate with maximum correlation and tracking delay in our dataset?

As a first step, we replicated the analysis conducted by Bonnen et al. (2015) by obtaining correlations between a) the log sensory noise and the tracking delay and b) the log sensory noise and the maximum correlation, separately for manual and ocular tracking. We used linear mixed models of the following structure:

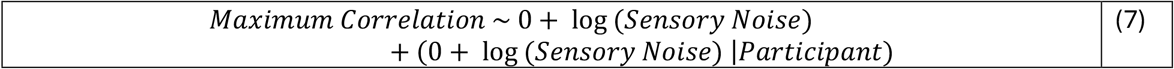

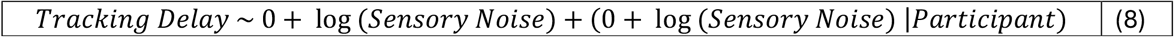

For manual tracking (Figure 9A and B), we found a regression coefficient of -0.14 (95% CI = [-0.16; - 0.12]; R^2 = 0.93) between sensory noise and maximum correlation and a regression coefficient of 0.143 (95% CI = [0.135; 0.15], R^2 = 0.91) for sensory noise and tracking delay. For ocular tracking (Figure 9C and D), the regression coefficients were -0.06 (95% CI = [-0.08; -0.05], R^2 = 0.83) between sensory noise and maximum correlation and 0.139 (95% CI = [0.131;0.146], R^2 = 0.86)) between sensory noise and tracking delay.

**Figure 9.**
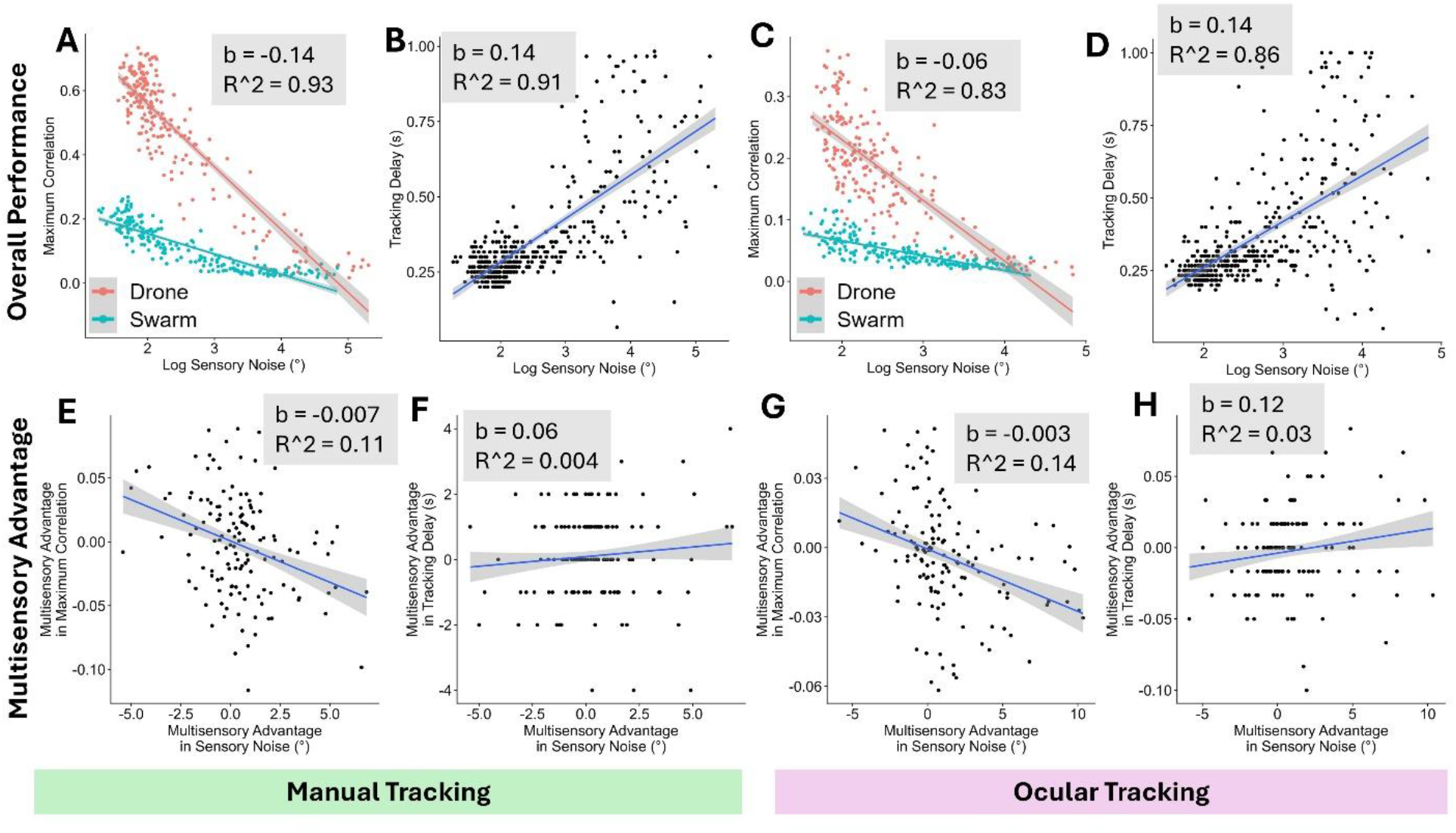
Correlations between the log sensory noise parameter and the maximum correlation (A and C) and between the log sensory noise parameter and the tracking delay (B and D), for manual tracking (A and B) and ocular tracking (C and D). E, F, G, and H: As A, B, C and D, but for the multisensory advantage in sensory noise, maximum correlation and tracking delay, respectively.

### Do these strong correlations hold for the multisensory advantage?

After establishing this baseline, we computed the multisensory advantage for each participant, fog level and tracking modality separately by subtracting the performance parameter in the no sound condition from the corresponding performance parameter in the sound condition:

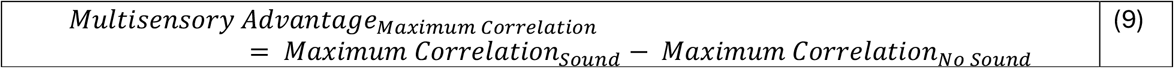

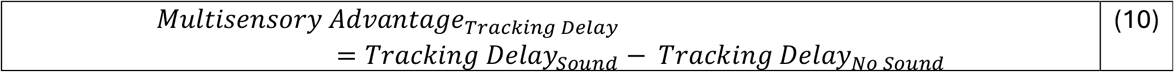

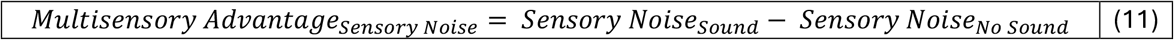

To avoid extreme values from biasing the estimates unduly, we removed all data points that fell more than 1.5 times the interquartile range above or below the upper/lower quartile. This was the case for between 5.5 and 13.4% of the data points depending on the condition; see Table 7 for a detailed breakdown.

**Table 7.** Number and percentage of excluded outliers for each multisensory advantage indicator.

| Dependent Variable –<br>Multisensory Advantage in... | # excluded | % excluded |
| --- | --- | --- |
| $\text{Maximum Correlation}_{\text{Manual}}$ | 13 | 5.5% |
| $\text{Maximum Correlation}_{\text{Ocular}}$ | 22 | 9.2% |
| $\text{Tracking Delay}_{\text{Manual}}$ | 19 | 8% |
| $\text{Tracking Delay}_{\text{Ocular}}$ | 32 | 13.4% |
| $\text{Sensory Noise}_{\text{Manual}}$ | 26 | 11% |
| $\text{Sensory Noise}_{\text{Ocular}}$ | 20 | 8.4% |

We then tested to what extent the multisensory advantage in the sensory noise correlated with the multisensory advantages in maximum correlation and tracking delay, respectively. Analogously to above, we fitted the following linear mixed models, separately for manual and ocular tracking.

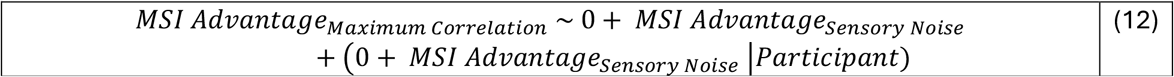

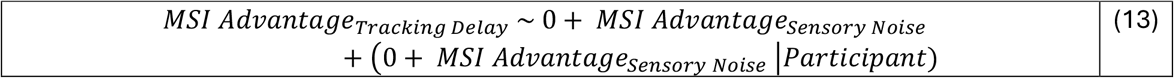

For the multisensory advantages observed during manual tracking (Figure 9E and F), we found regression coefficients of -0.007 (95% CI = [-0.012; -0.002], R^2 = 0.11) for sensory noise and maximum correlation and 0.06 (95% CI = [-0.08; 0.19], R^2 = 0.004). During ocular tracking (Figure 9G and H), the regression coefficients were -0.003 (95% = [-0.004; -0.002], R^2 = 0.11) for sensory noise and maximum correlation, and 0.12 (95% = [-0.06; 0.32], R^2 = 0.03) for sensory noise and tracking delay.

In agreement with Bonnen et al. (Bonnen et al., 2015), we found that the (log) sensory noise parameters explained a lot of the variability in the maximum correlations and tracking delay, both for manual and for ocular tracking (between 83% and 93%; see Figure 9A-D). This shared variability was much lower for the multisensory advantage, with values just above 10% for the maximum correlations (see Figure 9E and G). For the multisensory advantage in tracking delays, the multisensory advantage in the sensory noise explained barely any variability (see Figure 9F and H). This suggests that information gained by combining cues from different senses may not be used to guide behavior in the same way as visual information alone.

## Exploratory Analysis: The Relationship between Manual and Ocular Tracking

Another compelling feature of this dataset is that we obtained both manual and concurrent ocular response data for our participants in the context of a continuous psychophysics paradigm. This puts us in an ideal position to assess similarities and differences between the two response modalities. In this section we will therefore report exploratory analyses on the relationship between manual and ocular tracking more generally as well as on the relationship between sensory noise, tracking delay and maximum correlation specifically in the context of multisensory integration.

### How do Manual and Ocular Tracking Relate to each other?

We first examined the general relationship between manual and ocular tracking in our two experiments. For each dependent variable under investigation (maximum correlation, tracking delay, and sensory noise), we examined the correlation between values obtained from manual tracking and values obtained from ocular tracking. We fitted linear mixed models to the combined data sets of Experiment 1 and Experiment 2 with the ocular parameter (maximum correlation, tracking delay, sensory noise) as dependent variable, the corresponding manual parameter as fixed effect, and random slopes for the corresponding manual parameter per participant as random effects. We set both fixed effects and random effects intercepts to zero. The Wilkinson & Rogers notations of these three models read as follows:

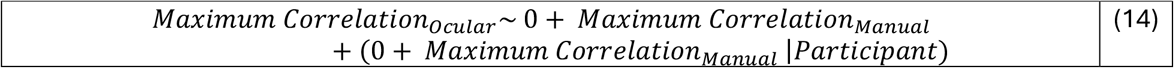

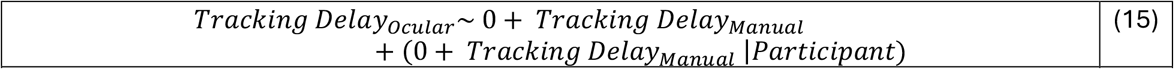

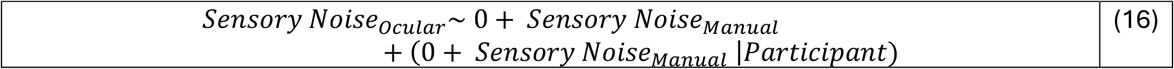

We computed bootstrapped 95% confidence intervals using the confint() function from base R. For the general performance (see Figure 10A-C), we found regression coefficients of 0.38 (95% CI = [0.36;0.41], R^2 = 0.92) for the maximum correlation, 0.95 (95% CI = [0.89; 1.01], R^2 = 0.009) for the tracking delays, and 0.78 (95% CI = [0.72;0.84], R^2 = 0.86) for the sensory noise.

**Figure 10.**
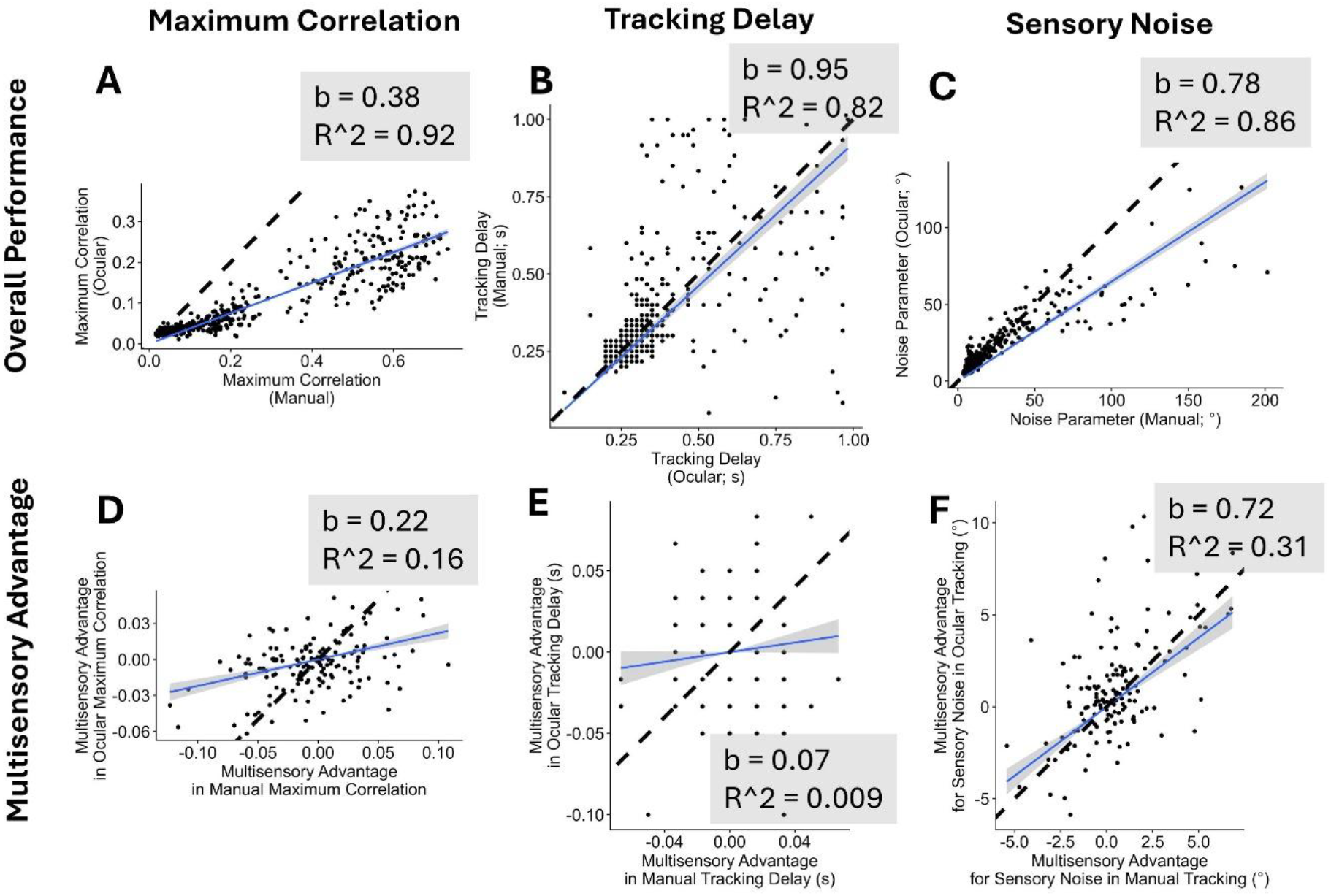
Correlations between manual tracking (x axis) and ocular tracking (y axis) for the maximum correlation (A), the tracking delay (B), the Kalman sensory noise parameter (C), the multisensory advantage in the maximum correlation (D), the multisensory advantage in the tracking delay (E) and the multisensory advantage in the Kalman sensory noise parameter (F). The blue line indicates a linear fit and the grey-shaded area is the 95% confidence band. The insets indicate the slope of this linear fit.

### Is added multisensory information used differently in manual and ocular tracking?

We then tested to what extent the multisensory advantage parameters correlated between manual and ocular tracking. Analogously to above, we fitted the following linear mixed models:

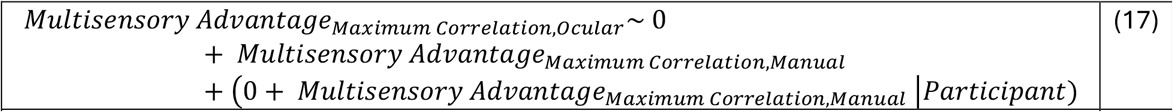

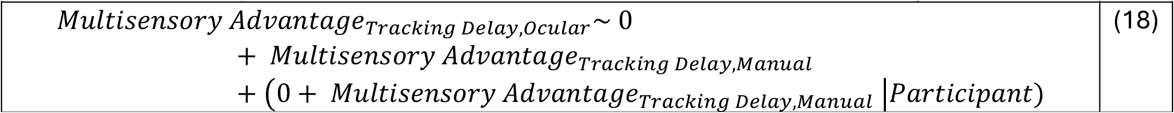

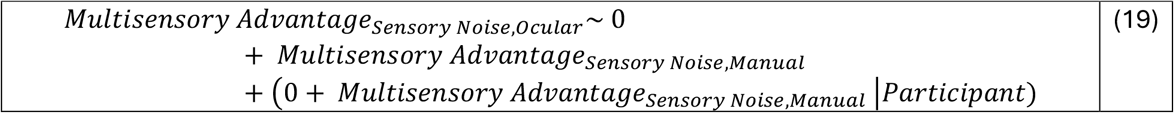

For the multisensory advantages (see Figure Figure 10D-F), we found regression coefficients of 0.22 (95% CI = [0.12; 0.32], R^2 = 0.16) for the maximum correlation, 0.07 (95% CI = [-0.19; 0.35], R^2 = 0.008) for the tracking delay, and 0.72 (95% CI = [0.54;0.96], R^2 = 0.31) for the sensory noise.

As expected, manual and ocular tracking were highly correlated across all three dependent variables with 82% to 92% shared variability. These correlations dropped substantially for the multisensory advantages with effectively no shared variability between manual and ocular tracking for the tracking delay.

## General Discussion

In this study, we set out to explore the use of continuous psychophysics in multisensory research, as instantiated in an audiovisual manual tracking task. As expected, we found that adding auditory cues to a visual stimulus enhanced sensory precision as measured by a Kalman filter-based analysis, particularly when visual cues were less reliable. This was reflected to some extent in the tracking variability (operationalized as the maximum correlation between stimulus and response), but much less reliably so than in the sensory precision. No multisensory advantages were found for the tracking delay (as measured by the time lag of the maximum correlation between stimulus and response). Simultaneous recording of eye movements revealed similar patterns for ocular tracking.

Our study provides some interesting insights into multisensory integration as measured by continuous tracking that complement Tonelli et al.’s (2025) findings. Tonelli et al. found no significant advantage for either of their dependent measures (tracking delay and maximum correlation) in their multisensory conditions over their audio-only or vision-only conditions, independent of the reliability of their visual stimulus. The data from our present study confirm this absence of a multisensory advantage in the tracking delay across both experiments, all fog conditions and both response modalities (see Figure 11B and E). The picture is, however, less clear for the tracking variability: we found a significant improvement in the maximum correlation for manual tracking under the highest visual uncertainty condition in Experiment 2 and for ocular tracking under the highest visual uncertainty in Experiment 1, and trends in the expected direction for some other conditions (Figure 11A and D). Given this mixed picture, it is our interpretation that a true effect is likely, i.e., multisensory integration does decrease tracking variability, but measurement noise makes it hard to detect. An important novel aspect of our study is the use of a Kalman filter to estimate participants’ sensory uncertainty underlying these performance measures: the sensory noise estimates show a much clearer multisensory advantage (see Figure 11C and F) that increases for fog conditions with higher visual uncertainty.

**Figure 11.**
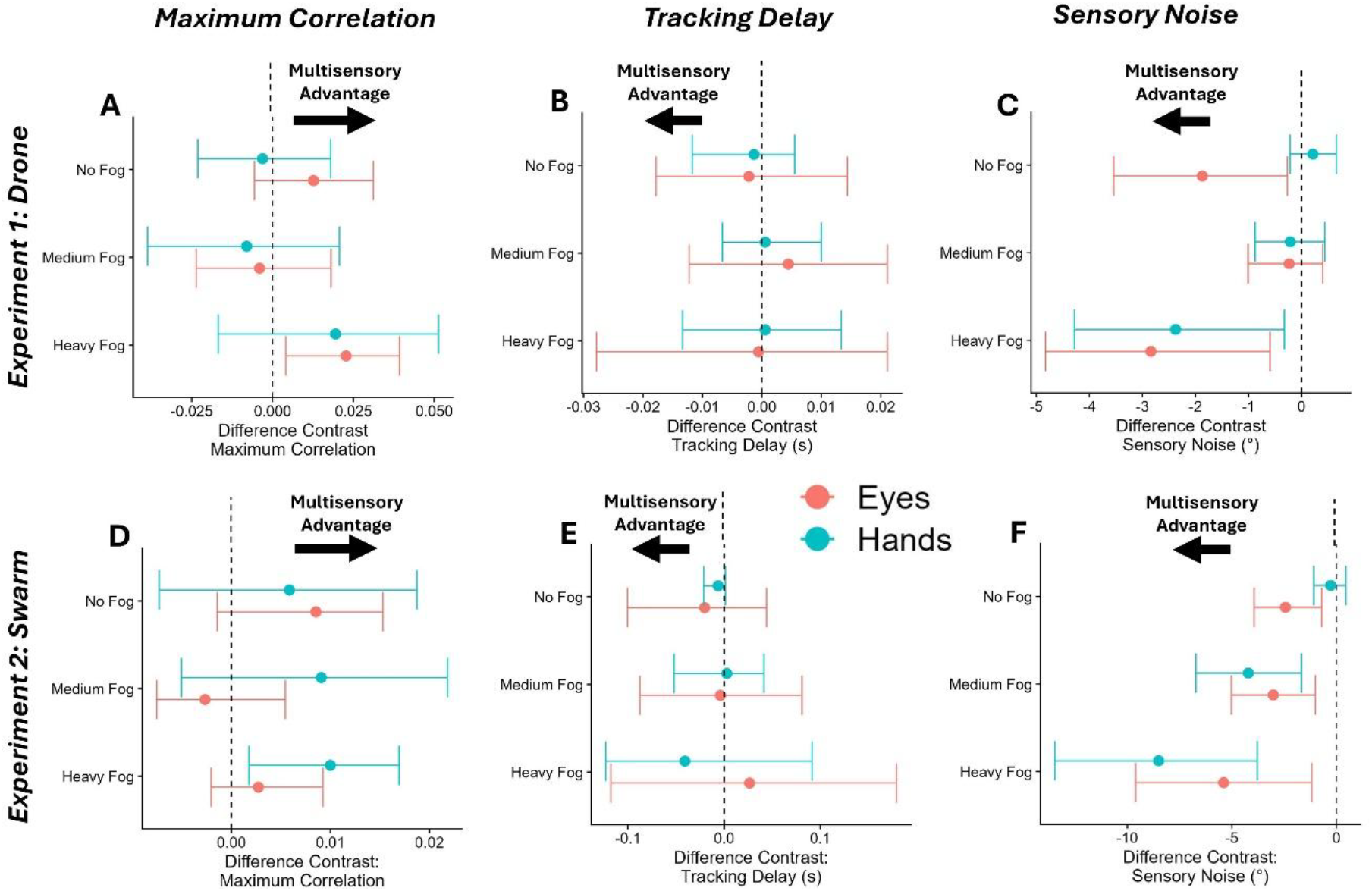
Regression coefficients and 95% confidence intervals (x axis) for the comparison between “sound” and “no sound” all three dependent variables (A and D: maximum correlation; B and E: tracking delay, C and F: sensory noise), both experiments (A, B, C: Experiment 1, D, E, F: Experiment 2), manual (blue) and ocular (red) tracking, and the three different fog levels (y axis). The dashed line indicates zero, i.e., no difference between the sound and the no sound conditions.

In exploratory analyses, we found – expectedly as per Bonnen et al. (2015) – high correlations between the fitted sensory noise parameters and the maximum correlations and tracking delays, respectively. However, when looking at the multisensory advantages in each of these dependent variables, i.e., how much higher maximum correlations, how much lower tracking delays, and how much lower sensory noise parameters were in the SOUND condition in comparison to the matching NO SOUND condition, the picture is much less clear: While there were still substantial correlations between sensory noise and the maximum correlations, the link between sensory noise and tracking delay was almost completely gone. Relatedly, the correlations in performance parameters between manual and ocular tracking were extremely high (with R^2s of 82% to 92%). However, when zooming in on the multisensory advantages, this correlation disappeared completely for the tracking delay and was substantially weakened for the maximum correlation and the sensory noise. These findings suggest that, in the study of visuo-auditory multisensory integration, these summary statistics (maximum correlation and tracking delay) may be insufficient proxies for the underlying sensory noise. This is further bolstered by our preregistered findings, where we found suggestive but inconsistent results for the maximum correlations and no multisensory advantage at all for the tracking delay; results were most consistent for the sensory noise parameter fitted using the Kalman filter.

## Conclusions

Overall, we were able to produce the increases in sensory precision for multimodal stimuli predicted by causal inference accounts of multisensory integration in a continuous tracking paradigm. This indicates that continuous psychophysics is generally suitable for the study of multisensory integration. However, in contrast to unimodal tasks, reliance on summary statistics alone may lead to an incomplete picture of the results, and modelling the data with a Kalman filter (Bonnen et al., 2015) or, as suggested more recently (Straub & Rothkopf, 2022), a Linear Quadratic Gaussian model, may be advisable.

### Open Science

You can download the Unity project as well as the executable, the data, the scripts used, and the pre-registrations for both experiments from Open Science Foundation (https://osf.io/6q7x3/).

We have deviated from the pre-registrations in the following ways:

- We implemented some improvements to the computational performance of our bootstrap analyses to reduce the overall runtime of the analysis
- There was an inconsistency between the manuscript and the code for the Kalman filter fitting procedure. The manuscript described negative loglikelihood minimization, while the preregistered code used an RMSE minimization. We changed the code to match the manuscript.
- The preregistration mentioned the plan to use to the data to build a version of the Drugovitsch et al. (Drugowitsch et al., 2014) model to accommodate continuous data. We dropped this plan because we deemed the data too noisy for this purpose. We changed the introduction from the pre-registration to reflect this change in framing and focus.
- The pre-registration did not account for the possibility that the equipment might fail to record eye or hand movements for substantial stretches of time. We excluded conditions where this occurred:
  - Experiment 1:
    ▪ No Fog + No Sound for participant d11
  - Experiment 2:
    ▪ Heavy Fog + Sound for participant e11
    ▪ Heavy Fog + No Sound, Heavy Fog + Sound, and Impenetrable Fog + Sound for participant e22
    ▪ Impenetrable Fog + Sound for participant e33
- We did not record our participants’ genders, and we are therefore unable to confirm that we met gender parity

